# Spatial extent of neighboring plants influences the strength of associational effects on mammal herbivory. Insights from a meta-analysis

**DOI:** 10.1101/019935

**Authors:** Emilie Champagne, Jean-Pierre Tremblay, Steeve D. Côté

## Abstract

There is high variability in the level of herbivory between individual plants from the same species with potential effects on population dynamics, community composition, and ecosystem structure and function. This variability can be partly explained by associational effects i.e. the impact of the presence of neighboring plants on the level of herbivory experienced by a focal plant, but it is still unclear how the spatial scale of plant neighborhood modulates foraging choice of herbivores; an inherently spatial process in itself. Using a meta-analysis, we investigated how spatial scale modifies associational effects on the susceptibility to browsing by herbivores with movement capacities similar to deer. From 2496 articles found in literature databases, we selected 46 studies providing a total of 168 differences of means in damage by herbivores or survival to woody plants (mostly) with and without neighboring plants. Spatial scales were reported as distance between plants or as plot size. We estimated the relationships between the effect sizes and spatial scale, type of associational effects and nature of the experiment using meta-analysis mixed models. The strength of associational effects declined with increasing plot size, regardless of the type of associational effects. Associational defences (i.e. decrease in herbivory for focal plants associated with unpalatable neighbors) had stronger magnitude than associational susceptibilities. The high remaining heterogeneity among studies suggests that untested factors modulate associational effects, such as nutritional quality of focal and neighboring plants, density of herbivores, timing of browsing, etc. Associational effects are already considered in multiple restoration contexts worldwide, but a better understanding of these relationships could improve their use in conservation, restoration and forest exploitation when browsing is a concern. This study is the first to investigate spatial patterns of associational effects across species and ecosystems, an issue that is essential to determine differential herbivory damages among plants.

## Introduction

Herbivory can modify the composition, structure and functions of ecosystems (Hester et al. 2006). There is high variability in the susceptibility of different plant species and individuals to herbivory. This variability is driven by forage selection, whom in itself is determined by the nutritional requirements of herbivores (Pyke et al. 1977), intrinsic (e.g. nutritive quality, Pyke et al. 1977), and extrinsic characteristics of both the plants and the environment (e.g. neighboring plants, Atsatt and O’Dowd 1976). Multiple studies have demonstrated the influence of neighboring plants on forage selection, a process named neighboring or associational effects (Milchunas and Noy‐Meir 2002, Barbosa et al. 2009), yet the conditions in which a specific neighborhood will increase or reduce herbivory are not fully understood. The distance between neighboring plants could explain part of the residual variability observed in associational effects (Underwood et al. 2014). Associational effects can be exploited as a management tool to alleviate the effect of herbivores; for example, Perea and Gil (2014) recommend planting seedlings under shrubs to reduce damage to the seedlings by browsers. Other recent studies (Noumi et al. 2015, Stutz et al. 2015, Torroba-Balmori et al. 2015) explored the application and limits of associational effects for the restoration of plant species, but without considering the spatial extent of plant neighborhood, although Stutz et al. (2015) quantified vegetation variables at two spatial scales. A better understanding of associational effects could improve and generalize their use in restoration, conservation and exploitation.

Four different types of associational effects have been described in the literature (Figure 1a), depending on the difference in palatability between the focal and the neighboring plants: (1) associational susceptibility involves a neighboring plant preferred to the focal plant, leading to increased consumption of the focal (Thomas 1986, Hjältén et al. 1993); (2) neighbor contrast defence describes the situation where the preferred neighbor concentrates the browsing pressure, thus decreasing herbivory on the focal plant (Bergvall et al. 2006, Rautio et al. 2012); (3) neighbor contrast susceptibility occurs when the less preferred or avoided neighbor leads to higher herbivory level on the focal plant (Bergvall et al. 2006; attractant-decoy hypothesis, Atsatt and O’Dowd 1976); (4) associational defence, or associational resistance, occurs when a less-preferred plant provides a protection from herbivory to the focal plant (Tahvanainen and Root 1972, Atsatt and O’Dowd 1976, Bergvall et al. 2006). A meta-analysis of all four associational effects by Barbosa et al. (2009) revealed that the direction and strength of effects are influenced by herbivore taxonomy (e.g. mammals or insects), plant taxonomic relatedness and the palatability of the neighboring plant, but unexplained variation remains. The focus of this meta-analysis is the contribution of the spatial scale of the neighborhood to the unexplained variation in associational effects.

**Figure 1.**
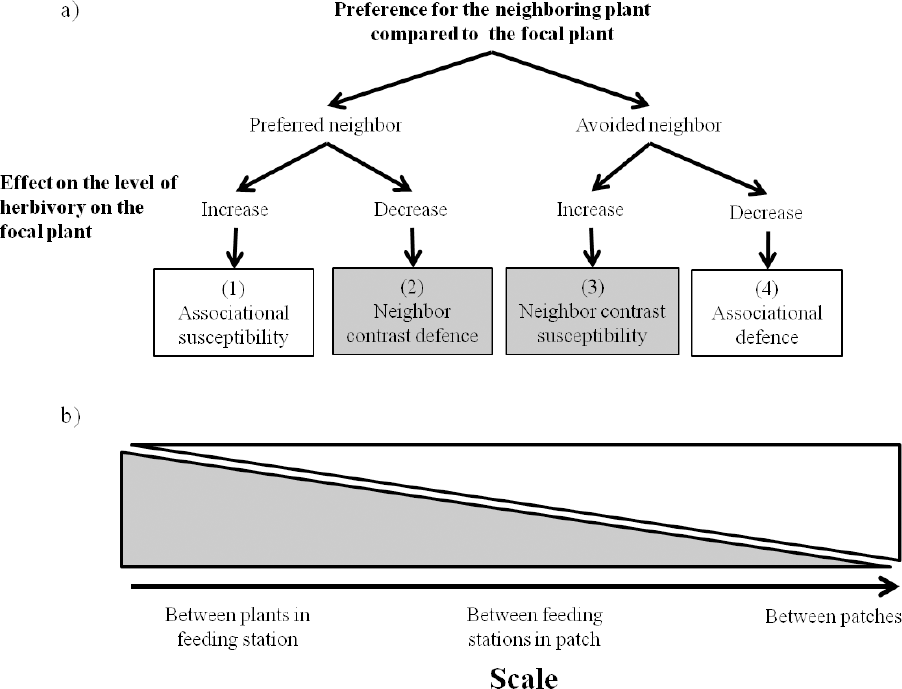
a) Flowchart of the type of associational effects affecting the level of herbivory on the focal plant based on the preference of the herbivore for the neighboring plants versus the focal plant (first level of the flowchart) and on the direction of the association (second level). “Classic” types of effects (associational susceptibility and defence) are in white boxes while “contrast” types (neighbor contrast susceptibility and defence) are in grey boxes. b) Predictions about how the “classic” (white) and “contrast” (grey) associational effects should vary in strength with spatial scale according to Bergvall et al. (2006) framework. Scales suggested on the x-axes are suggestions not representing exactly where the type of associational effects are expected to occur.

Forage selection is an inherently spatial phenomenon and its impacts can be measured at multiple spatial scales from the choice of a single bite to the establishment of a home range within the distribution range of a population (Johnson 1980, Brown and Allen 1989, Bommarco and Banks 2003). At the scale of the feeding site or the patch, Bergvall et al. (2006) predicted higher occurrence of associational susceptibility and associational defence effects (Figure 1, effects 1 and 4). The decision to use a patch should be a function of the relative attraction of adjacent patches based on the palatability and abundance of plants composing them favoring classic susceptibility or defence effects (Figure 1, effects 1 and 4). Within a patch, Bergvall et al. (2006) predicted higher occurrence of neighbor contrast defence or susceptibility (Figure 1, effects 2 and 3), because the choice made by the animal would then be a function of its ability to detect differences in palatability of adjacent plants. Although multiple spatial scales have been tested with invertebrate herbivores (Thomas 1986, Karban et al. 2006, Karban 2010), few experiments have tested the effect of hierarchical foraging on associational effects. Exceptions include a study of red deer (*Cervus elaphus*) and sheep (*Ovis aries*) showing decreased herbivory on *Calluna vulgaris* with increasing distance from preferred grass patches (Hester and Baillie 1998); this associational susceptibility disappeared at 1 to 3 m from the grass patch, depending on herbivory pressure. Bergvall et al. (2006) tested the selection of fallow deer (*Dama dama*) between patches and within patches of pellets with varying tannin concentration. They found that palatable food was consumed more in the immediate neighborhood of highly defended food (neighbor contrast susceptibility) and highly defended food was consumed less in a high palatability neighborhood (neighbor contrast defence). Underwood et al. (2014), also raised that empirical studies and modeling of associational effects currently lack consideration for the role of spatial scale.

Here, we used a meta-analysis approach to determine whether the spatial scale modulates associational effects of neighboring plants on the level of herbivory. Because dispersal can affect the potential for large scale associational effects (Grez and Gonzalez 1995), we controlled for differences in dispersal capacity by restricting our study to herbivores with movement capacities similar to deer, i.e. from small deer such as roe deer (*Capreolus capreolus*) to moose (*Alces alces)*, and including herbivores from other groups of similar body sizes, such as wild boar (*Sus scrofa*) and Western grey kangaroo (*Macropus fuliginosus*). Our first objective was to characterize how associational effects vary in strength, depending on their type (numbers 1 to 4, Figure 1). Second, we described how associational effects vary in strength with the spatial scale considered. We hypothesized that hierarchical forage selection determines the most frequent type of associational effects within and between patches, i.e. the “classic” type (associational susceptibility and associational defence) or the “contrast” type (neighbor contrast defence and susceptibility), according to the conceptual framework provided by Bergvall et al. (2006). We thus predicted an interaction between distance and associational effect type (Figure 1b) where associational susceptibility or defence would be more frequent at larger spatial scales (home ranges, patches) when herbivore select resources based on the relative abundance of resources, while “neighbor contrast” would be more frequent once herbivores are feeding within a patch and selecting individual plant species. This study is the first to investigate how spatial scale drives associational effects across herbivore species and ecosystems, an issue essential for understanding variations in the level of herbivory incurred by individuals within a population (Barbosa et al. 2009, Underwood et al. 2014).

## Methods

### Literature review

We obtained 2496 peer-reviewed publications using the search strategy presented in Appendix A in ISI Web of Science, Biosis preview and BioOne (in July 2013), and through citations found in these publications. We searched for studies involving herbivores with movement capacities similar to deer from the smallest to the largest deer species; the smallest herbivore in our dataset is European roe deer and the largest is the European bison (*Bison bonasus)*. Studies reported data on damage or survival of plants (hereafter called the focal plants) with and without the presence of a neighboring plant (hereafter called the neighbor plant). Damage was inferred from counts of browsed twigs or leaves, or biomass removal and did not include measures of growth or regrowth following herbivory. We included studies using feeding trials in controlled or natural environments, transplantation/removal of neighbors and observations in natural environments.

We established the criteria regarding acceptance or rejection of a study prior to conducting the meta-analysis using a PRISMA inspired protocol (see process in Appendix A, Moher et al. 2009). The criteria were the presence of a control treatment (herbivory without neighboring plant), a palatable plant in the focal-neighbor group, and a difference in palatability between plants. To evaluate the effect of spatial scale, each study needed to clearly state the size of the plot where data were recorded or the distance between the focal and neighboring plant. We rejected data on seed predation a posteriori. A single observer (EC) reviewed and selected all articles and recorded each rejection criterion. To ensure the reproducibility of study selection, a second observer screened a subsample of 460 publications; the first and second observers agreed on 456 publications (452 rejected, 4 accepted) leading to a kappa statistic (Cohen 1960) of 0.66, exceeding the level of 0.60 and thus indicating that publication selection was reproducible (Côté et al. 2013). Following this procedure (Appendix A), we kept 46 publications from the original 2496 (Supplement).

### Data extraction and effect size computation

For each article, a single observer (EC) extracted information regarding the study, such as the nature of the experiment, identity of the herbivore, plot size, etc. (see Appendix B for a complete list). To compare associational effects among studies, we extracted means and variance of damage and/or survival with and without neighboring plants. We used this information to compile standardized effect sizes that indicate the size of the impact of a neighboring plant on herbivory on the focal plants (see below for details). We also extracted independent variables, such as the type of associational effect (“classic” or “contrast”, Figure 1) and the direction of the effect. By direction, we mean the effect on the level of herbivory on the focal plant (Figure 1), which is increase in herbivory (now referred as the susceptibility subgroup) or decrease in herbivory (now referred as the defence subgroup). Some studies measured associational effects in plots while others reported a linear distance between focal and neighbor plants. We decided not to combine the plot-based and distance-based studies because of the variation in the spatial range they covered (plot-based studies: range varying from 0.01 m^2^ to 148 000 m^2^ with a median = 27.5 m^2^, distance-based studies: range from 0 to 2 m, median = 0.02 m). Focal plants located underneath their neighbor without further indication were given a distance value of 0. Variables extracted from articles are detailed in the Appendix B. Data presented in graphs were extracted using Web Plot Digitizer V2.5 (Copyright 2010-2012 Ankit Rohatgi). We contacted authors for missing data, such as plot size, variance, Pearson’s r or identity of the herbivore species (See supplementary Table 2).

The data extraction provided 283 distinct observations of damage/survival with and without neighboring plants. Data reported as means with variance were transformed into standardized mean difference (*d*), a common effect size used for meta-analysis in ecology (Borenstein et al. 2009, Rosenberg et al. 2013). In the few cases where data were reported as percentage of all focal plants browsed, we computed log odd ratios (OR) using a 2 x 2 contingency table with browsed/unbrowsed columns and with/without neighbors rows (Borenstein et al. 2009, Rosenberg et al. 2013). Other studies correlated damage to the abundance (e.g. cover) of the neighbor species and reported Pearson’s r as an effect size statistic (Borenstein et al. 2009, Rosenberg et al. 2013). Depending on whether the direction of the effect was susceptibility or defence, values of *d* and Pearson’s r could be negative or positive. We transformed them into absolute values as the categorical variable “direction” already reports whether they belong to the increased susceptibility or increased defence subgroups (Appendix B). Effect sizes computed as OR and r were converted into *d* and added into a single analysis using equations from Borenstein et al. (2009). We selected *d* for common effect size as most data were available as a difference of means (Appendix B) and because of its simple interpretation; the higher the *d* value, the greater is the influence of the neighboring plant on the focal plant herbivory level. Although not frequently used (but see Hamm et al. 2010, Thomson et al. 2013), converting effect sizes allows the inclusion of all data answering the same broad question and avoids information loss through rejection of relevant studies (Borenstein et al. 2009).

When confronted with multiple effect sizes from one study, we extracted them all, unless a global mean was available (e.g. Russell and Fowler 2004). In the final analysis, we kept only one combination of neighboring plants, herbivore and spatial scale (distance between neighbors or plot size), similar to Barbosa et al. (2009), which meant keeping more than one effect size per study in some cases. When the same combination occurred in the same study, we combined those redundant effect sizes following Borenstein et al. (2009) (Appendix A and Supplement for details). Following those steps, we obtained a total of 168 effect sizes from 44 studies.

### Statistical analyses

We tested the impact of independent variables on the standardized difference of mean (*d*) in three meta-analysis mixed models using the function *rma* of the metafor package (Viechtbauer 2010) in R 3.1.2 (R Core Team 2013). For our first objective, we used the complete dataset to test the variation in effect size depending on the direction of the association (susceptibility, defence; figure 1a), type of association (“classic”: associational defence/associational susceptibility, “contrast”: neighbor contrast defence/neighbor contrast susceptibility; figure 1a) and interaction between direction and type of association. We also included the nature of the experiment (feeding trial, observation study, transplantation or removal experiments) since effect sizes from controlled experiments such as feeding trials could be stronger than results of observational studies where foraging by herbivores would be influenced by uncontrolled factors. The conversion of OR and r in *d* could have generated a bias in the values of the effect sizes; we tested this supposition in a simple model with effect size class (d, r or OR) as an independent variable. Since effect size class did not influence the value of *d* (*d*-class compared to OR-class: z = -0.2, p = 0.8; compared to r-class: z = -0.5, p = 0.6), we did not include it in our final model.

For our second objective, we tested the effect of spatial scale on associational effect strength for plot-based and distance-based studies separately. We log-transformed plot size to control for its large dispersion (Bland and Altman 1996). For both models, together with the variables describing the linear and quadratic parameters for the spatial scale (log plot size or linear distance), we included the type of association and their interactions to test for predictions of higher frequency of “classic” interaction at a finer scales and higher frequency of “contrast” interaction at a larger scales (Figure 1b). Both models also included the nature of the experiment as an independent variable to control for differences in effect sizes from different experiments.

The function *rma* weights effect sizes using the inverse-variance method for mixed models following this equation (Viechtbauer 2010):

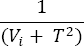

where V_i_ is an estimate of the within-study variance and *T*^2^ an estimate of between-study variance, calculated from the effect sizes. The percentage of heterogeneity in the effect sizes explained by independent variables was estimated by how much the addition of variables reduced the estimate of residual heterogeneity (Viechtbauer 2010). We further evaluated the heterogeneity of all effect sizes inside each level of independent variables by calculating the value of I^2^, the proportion of observed variance reflecting real differences among effect sizes (Borenstein et al. 2009); a 0 value of I^2^ indicates no between-study variation, while a high value indicates untested independent variables.

We tested the sensibility of our model to outliers (Viechtbauer and Cheung 2010) using the function *influence* of the metafor package (Viechtbauer 2010). We tested for publication bias using funnel plots with Egger’s regression test (Sterne et al. 2001, Jennions et al. 2013) and the trim and fill method (Duval 2005, Jennions et al. 2013), using the *regtest* and *trimfill* functions of the metafor package for R 3.1.2 (R Core Team 2013) with standard error as the predictor (Viechtbauer 2010). Additionally, we performed a cumulative meta-analysis and tested year of publication as an independent variable to ensure the absence of a temporal trend in the effect sizes (Koricheva et al. 2013). All statistical analyses were performed using α = 0.05 and results are presented as means with 95% confidence intervals.

## Results

The selected studies reported results related to over 51 focal plant species; 15 were reported in more than one article and only one out of 15 was not a woody plant (*Medicago sativa*). Most woody plants were reported in two to three studies, *Pinus sylvestris* and *Picea abies* were the focal species in 11 and six articles, respectively. Over 70 different neighbor plant species were found; *Betula pendula* was present in five articles, but most neighbor species were reported in only one study. Twelve studies reported domestic sheep (*Ovis aries*) as the main herbivore. *Alces alces* and *Capreolus capreolus* were mentioned in eight studies and *Cervus elaphus* in seven studies. The extracted data were equally distributed between decreased and increased herbivory with neighboring plant, but “classical” types (associational defence and associational susceptibility, n = 104) were more frequent than “contrast” types (neighbor contrast defence and neighbor contrast susceptibility, n = 47). Most effect sizes resulted from feeding trials (n = 71), where various assemblages were proposed to herbivores, but 54 came from observational studies and 38 from transplantation experiments. Removal experiments were rarely used (n = 5). Additional summary data can be found in Appendix B.

The first model using the complete dataset explained 23% of the heterogeneity between effect sizes (omnibus test for independent variables: Q_df = 8_ = 50.0, p < 0.0001) and the pseudo-R^2^ for the model reached 23.0%. There was, however, a high residual heterogeneity in the model (test for residual heterogeneity: Q_df = 159_ = 1047.0, p <0.0001). Effect sizes for defence associational effects (associational defence and neighbor contrast defence) had a greater magnitude than susceptibility associational effects (associational susceptibility and neighbor contrast susceptibility; Figure 2). Classic associational effects also had a greater value than contrast associational effects (Figure 2). Except for the contrast level of associational effects, all I^2^ were above 70%, indicating the presence of untested variables (Figure 2). Transplantation experiments presented the strongest and more variable values of *d*, while feeding trials found consistently small associational effects (Figure 2); values for observational studies were intermediate (Figure 2).

**Figure 2.**
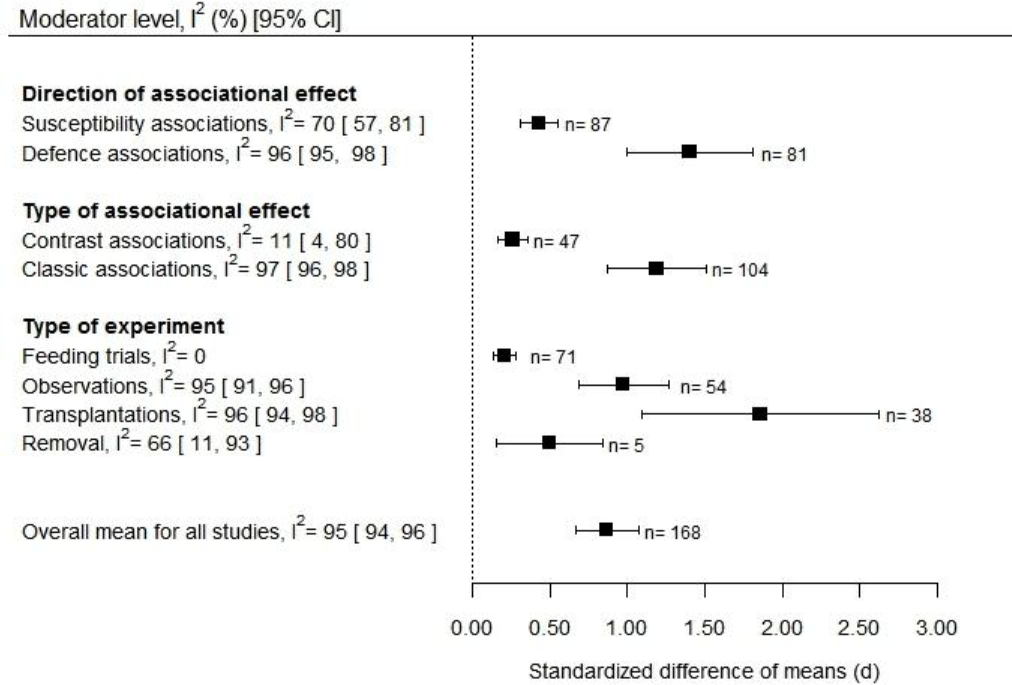
Summary of difference in damage/survival with and without a neighboring plant (*d*, standardized difference of means) separated by the independent variable levels tested, with 95% CI and I^2^, the percentage of total variability due to heterogeneity among *d*’s. A higher *d* indicates a higher associational effect of the neighboring plant on the focal plant’s herbivory level. Numbers to the right of the data points are the number of effect sizes in each summary effect. We used a meta-analysis mixed model to test the impact of variables on the standardized difference of means.

The model of the effect of plot size on associational effects explained 68% of the heterogeneity (omnibus test for independent variables Q_df= 9_ = 28.5, p = 0.0008, pseudo-R^2^ = 19.6 %) but also presented high remaining heterogeneity (Q_df = 86_ = 312.9, p <0.0001). As the log-plot size increased, there was a linear decrease in the strength of associational effects (Figure 3a, estimate = -0.13 [-0.22, -0.05]). There was no interaction between the type of associational effect and plot size (z = -0.22, p = 0.8). The model of the relationships between associational effect size and distance between the focal and neighboring plant explained a low amount of heterogeneity (3%; pseudo-R^2^ = 19.1 %; omnibus test for independent variables Q_df= 6_ = 20.5, p = 0.002) and consequently had a high amount of remaining heterogeneity (Q_df = 65_ = 674.0, p <0.0001). There was no effect of the distance between neighbors on the strength of associational effects (linear estimate: z = -0.1, p = 0.2; quadratic estimate: z = -0.1, p = 0.9), nor of the interaction between distance and type of associational effect (z = 0.4, p = 0.7). Visual examination of the data revealed a sharp decline in effect size after 0.1 m (Figure 3b).

**Figure 3.**
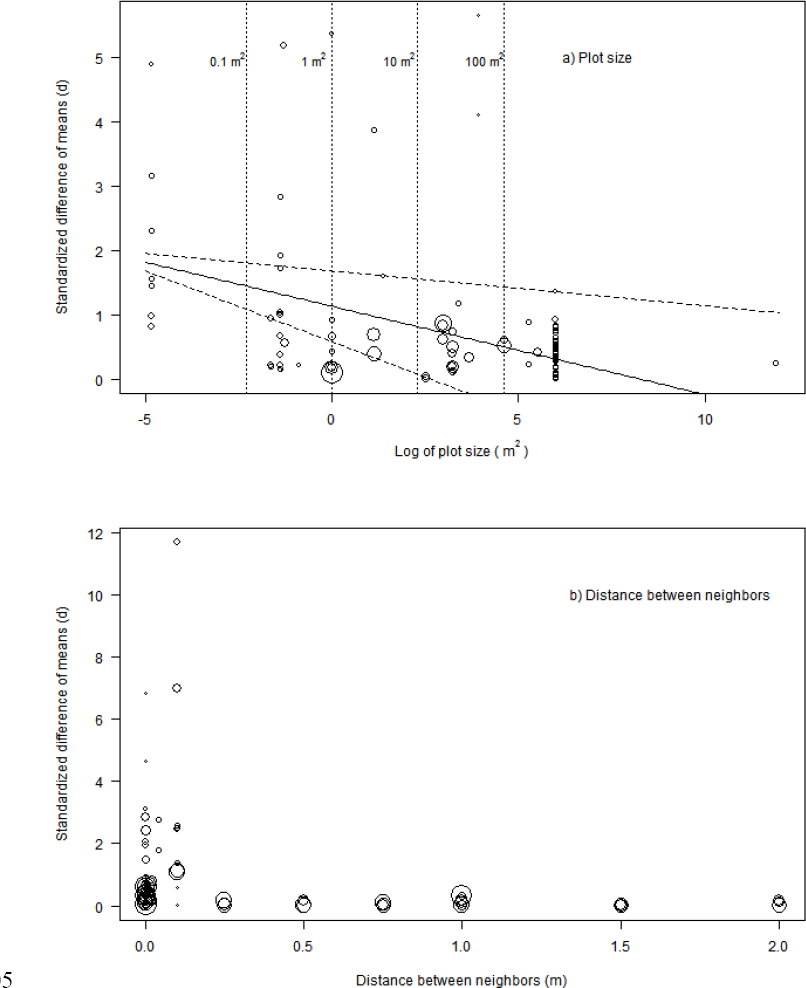
Relationship between associational effects and two different indicators of spatial scale: a) decrease in the difference in damage/survival with and without a neighboring plant (standardized difference of means) according to plot size (m^2^); b) associational effect limited to the first 10 cm between the focal plant and its neighbors. For each figure, the size of each point indicates the weight of each effect size in the meta-analysis mixed effect model, calculated with the inverse-variance method. Vertical lines and numbers above correspond to untransformed values of plot size (m^2^). Regression line results from a meta-analysis mixed model and dotted lines represent predicted values with 95% CI.

The sensitivity analysis for outliers uncovered nine effect sizes that could potentially change the results in the associational effect type model, seven in the spatial scale model with plot-based studies and three in the spatial scale model with distance-based studies. We analyzed each of the models without each of their outliers sequentially. In the associational effect type model, the removal of the data from a transplantation study (ID 156-157, Supplement) makes the nature of the experiment different (observational studies significantly higher from the others) while removing ID 64 (Häsler and Senn 2012) generates an interaction between type and direction of effect size. The effect size from that study was computed from two particularly high R^2^ values (0.96 and 0.61), combined as they represented a single combination of plants, distance and herbivores. Removing the only observation presenting a very large spatial scale (size = 148 000 m^2^, DeGabriel et al. 2011) did not modify the relationship with plot size in the spatial scale model. Because there was no reason to exclude any of those effect sizes based on the study characteristics, we kept the outliers in the final model (Viechtbauer and Cheung 2010). We also found some evidence of potential publication bias in funnel plots for the entire dataset and used the trim and fill method to test the robustness of the overall mean effect size (Appendix C). The trim and fill method identifies and correct the asymmetry by imputing smaller effect sizes around an estimated true center (Viechtbauer 2010). For the entire dataset, the trim and fill method generated more values of associational susceptibilities, suggesting either a publication bias in the analyses or a naturally higher frequency of associational defences (Appendix C). In addition, our analyses revealed potential publication bias among the effect sizes calculated as difference of the means (effect size of class *d*) and in observational studies (Appendix C). Even with input values, the *d*-class subgroup mean is similar to the r and OR-class subgroups and thus should not modify our conclusions. The trim and fill method suggests more associational susceptibilities in the observational studies subgroup, but this asymmetry could also result from the higher natural occurrence of associational defences. We found no evidence of a temporal trend (Appendix C).

## Discussion

Using a meta-analysis based on 46 studies and 168 data points on associational effects of neighboring plants on the level of herbivory, we found a decrease in associational effect strength with spatial scale. In contradiction with our hypothesis, the decrease was independent of the type of associational effect (i.e. “classic” or “contrast” type). We also found that associational defences had stronger effects than associational susceptibilities. There is a common agreement that hierarchical forage selection has been overlooked in associational effect studies (Barbosa et al. 2009, Hambäck et al. 2014, Underwood et al. 2014). Our study is the first pointing out the magnitude of change in associational effects with spatial scale.

The descriptors of spatial scale, i.e. presence of neighbors in a plot or distance between focal and neighbor, highly influenced the relation between scale and associational effects. Distance between plants is a one-dimensional measure, mostly used when studying the relationships between two individual plants (e.g. nurse plant studies or in feeding trials). This is reflected by the small range of distances in our dataset. When considering those simple interactions, associational effects declined quickly with increased distance between the plants. Typical mechanisms of associational effects, like reduction of apparency of the focal plant or induction of chemical defence (Barbosa et al. 2009), could only be expected when neighboring plants are close to one another. On the other side, multiple focal and neighboring plants can be present in a plot, complexifying the interactions, thus possibly explaining the slower decline of associational effects with increasing scale. Resource selection and enerfy maximization by herbivores could also explain large scale associational effects (Courant and Fortin 2010). Even if the strength of associational effects decreases with plot area, a predicted *d* of 0.82 for 10 m^2^ plots is still a large effect size according to Cohen’s rule of thumb (Cohen 1988). Experiments with relatively large plots (196 m^2^, Danell et al. 1991; 400 m^2^, Milligan and Koricheva 2013 and Vehviläinen and Koricheva 2006) also presented large *d* according to Cohen (1988). The information reported in the publications prevented us from testing the effect of the relative density between focal and neighboring plants, but this would probably explain part of the variation in associational effects in larger plots. Few studies investigated associational effects at large distance or in very large plots. Moore et al. (2015) recently demonstrated associational susceptibility and neighbor contrast defence for *Calluna vulgaris* within 1000 m of grass patches in the Scottish heathlands.

We did not find support for the predictions that “classic” effects should influence patch choice by herbivores while “contrast” effects should affect within patch selection (Bergvall et al 2006). Because few associational effects reported were measured in large patches, the model could have been unable to detect an interaction between type of association and distance. Every type of effects could also be seen at all scales because of the additive effects of herbivore selection at multiple scales (Miller et al. 2006). The associational effect seen at a specific scale could result from the addition of associational effects at other scales; fine scale associational susceptibilities or defences could be triggered by large-scale distribution of neighboring plants. This could be particularly important in studies performed in natural environments. Aside from Bergvall et al. (2006) and their following work (Bergvall et al. 2008, Rautio et al. 2008, Rautio et al. 2012), few authors have studied how spatial scaling relates to associational effects through the foraging behavior of large herbivores (but see Courant and Fortin 2010, Wang et al. 2010, Stutz et al. 2015). For small mammals, Emerson et al. (2012) tested associational effects at three spatial scales (between patches > between feeding stations > within feeding stations) with squirrels (*Sciurus* spp.), and found that both neighbor contrast susceptibility and associational defence occur among patches and among feeding stations. At a larger scale, they found only associational defence; high palatability seeds were less susceptible to be consumed in low palatability patches. The study of associational effects could be greatly improved by more experimentation with varying patch size and distance between neighbors, which could test the extent of associational susceptibilities and defences such as the study by Oom and Hester (1999).

Associational defences had stronger effects than associational susceptibilities, thereby suggesting stronger effects of facilitation. Facilitation between plants is known to be common in stressful environments, such as those with high herbivory pressure (Callaway and Walker 1997). High herbivory pressure, however, can also reduce the impact of associational defences, as herbivores could become less selective when competition between individuals increases (Baraza et al. 2006). Some studies have demonstrated a relation between herbivory pressure and associational effects (Aerts et al. 2007, Graff et al. 2007, Smit et al. 2007), but the heterogeneity in reporting herbivore pressure prevented us to test this factor. “Classic” type of associational effects also presented stronger effects than “contrast” type. Although Atsatt and O’Dowd (1976) introduced the attractant-decoy hypothesis 40 years ago, interest in contrast associational effects is more recent (see Bergvall et al. 2006) and they might be understudied; only 47 of our effect sizes concerned “contrast” interactions.

The strength of associational effects were also dependent on the nature of the experimental design. We expected observational studies to have low and variable associational effects, since the environment is uncontrolled and thus more variable. Surprisingly, feeding trials reported the lowest associational effect sizes, and transplantation experiments in natural environments reported effects of the highest magnitude. The simplicity of the feeding trials could explain the low values and low variance of those associational effects. As demonstrated by Wang et al. (2010), complex neighborhood can provide associational defence, either by a passive reduction of selectivity or by generating mistakes in foraging choices. They reported that the palatable grass *Medicago sativa* was less consumed by sheep in complex heterogenous environment including three plant species compared to homogenous environment (Wang et al. 2010). Herbivores integrate information at multiple spatial and temporal scales in natural environments to make foraging decisions (Miller et al. 2006) thereby generating associational effects.

In their meta-analysis, Barbosa et al. (2009) stated that associational defence was the most frequent associational effect under mammalian herbivory. Our results indicate, however, that associations with a plant providing defence (n = 81) are not more frequent than associations with a plant increasing consumption (n = 87). The asymmetry found in effect sizes could be an indication that associational defences are more frequent as the distribution of effect sizes is skewed towards them, but could also result from publication bias. Our dataset is dominated by woody plants already including a large variation in functional traits, still consideration for a wider range of functional types could help disentangle which of increased defence or susceptibility in presence of neighbors is more prevalent for herbivores with movement abilities similar to deer. Woody plants could be more apparent to herbivores than herbaceous plants because of their larger size and longer life span (Haukioja and Koricheva 2000) and those differences could be reflected in associational effects. Most studies of associational effects involving herbaceous species that we reviewed measured parameters such as growth, height or survival of individuals that did not always allow distinction of the effects of herbivory from interactions such as competition or facilitation.

As with many meta-analyses, there are restrictions to the generalization of our results. First, our work focused on herbivores with movement abilities similar to deer and the results cannot be exported to smaller mammals or invertebrates, as their foraging behavior is much different. Small herbivores are relatively more selective than larger ones and can discriminate between plants and plant parts at finer spatial scales so we should not expect associational effects of the same magnitude (Olff et al. 1999). For example, in one study roe deer (*Capreolus capreolus*) selected forages at both patch and plant levels, while rabbits (*Oryctolagus cuniculus*) selected plants only at the species level and were not influenced by the spatial arrangement of plants (Bergman et al. 2005). Second, the large heterogeneity found in effect sizes (Figure 2) indicates that many untested variables influenced the magnitude of associational effects and their interactions with scale. For example, we did not take into account the season; in seasonal environments, selectivity could be lower in winter because of the lack of resources, or higher given energy constraints, respectively reducing or increasing the strength of associational effects. Many of the studies included in our meta-analysis presented survival or damage for an entire year and we combined the data from multiple seasons or years, which partly explain the remaining heterogeneity. Our goal was to explore general patterns, but we contend that multiple factors can influence associational effects, such as relative abundance or density of focal or neighbor plants (Emerson et al. 2012, Hambäck et al. 2014, Underwood et al. 2014), richness of food patches (Milligan and Koricheva 2013), diversity (Castagneyrol et al. 2014), herbivore density (Aerts et al. 2007, Graff et al. 2007, Smit et al. 2007), etc. Finally, our sensitivity analyses for outliers and recombined effect sizes showed a consistent negative effect of plot size on the value of effect sizes.

Our study updates and extends previous work, providing new insights that should fuel further research, on the spatial range of associational effects, the spread of contrast type interactions and the prevalence of associational defence and susceptibility in large herbivores. We suggest a more systematic reporting of contextual data, such as herbivore densities, herbivores diet breath and densities of neighboring and focal plants, as those variables could explain the high residual heterogeneity of associational effects.

## Acknowledgements

This project is part of the Natural Sciences and Engineering Research Council of Canada (NSERC) Chair in integrated resource management of Anticosti Island (http://www.chaireanticosti.ulaval.ca/). E.C. received scholarships from NSERC. We thank all the researchers who shared data of their study, M. Courchesne for help with article selection steps, and A. Hester, V. Saucier, P. Morissette and M. Bonin for reviewing an earlier version of this manuscript. H. Crépeau from the Service de consultation statistique at Université Laval provided statistical guidance. Two anonymous reviewers provided valuable comments to improve an earlier version of this manuscript.

